# Characterising and Minimising Step and Filtering Artifacts in TMS-EEG Recordings

**DOI:** 10.64898/2026.05.13.725016

**Authors:** Marissa M. Holden, Mitchell R. Goldsworthy, Wei-Yeh Liao, Scott R. Clark, Christopher C. Cline, Corey J. Keller, Julio C. Hernandez-Pavon, Nigel C. Rogasch

**Affiliations:** School of Pharmacy and Biomedical Sciences, College of Health, Adelaide University; Hopwood Centre for Neurobiology, Lifelong Health Theme, South Australian Health and Medical Research Institute; School of Psychology, College of Education, Behavioural and Social Sciences, Adelaide University; School of Medicine, College of Health, Adelaide University; Department of Psychiatry and Behavioral Sciences, Stanford University, Stanford, CA, USA; Wu Tsai Neurosciences Institute, Stanford University, Stanford, CA, USA; Veterans Affairs Palo Alto Health Care System, Palo Alto, CA, USA; Department of Psychological Sciences, Kansas State University, Manhattan, KS, USA; School of Psychological Sciences and Turner Institute for Brain and Mental Health, Monash University, Australia

**Keywords:** Transcranial magnetic stimulation (TMS), Electroencephalography (EEG), TMS-evoked potentials (TEPs), Artifacts, Filtering

## Abstract

Transcranial magnetic stimulation combined with electroencephalography (TMS-EEG) enables direct measurement of cortical reactivity via TMS-evoked potentials (TEPs). Interpretation of early TEP components however, is highly sensitive to stimulation and hardware-related artifacts. We identified and characterised a persistent, non-neural ‘step-drift’ artifact unexpectedly present in recent TMS-EEG recordings from our group. We show that the artifact is distinct from previously described TMS pulse and discharge/decay artifacts and likely reflects a hardware interaction phenomenon. We demonstrated that amplifier settings, but not TMS pulse shape, substantially influenced artifact expression, with DC-coupled recordings with no online high-pass filter reducing step amplitude compared with AC-coupled recordings with a high-pass filter. Simulations additionally revealed that filtering over the step-drift artifact introduced pronounced ringing and edge artifacts, highlighting the need to address this artifact prior to data processing. We propose a processing pipeline incorporating robust polynomial detrending and a modified Butterworth filter with autoregressive extrapolation that minimised TEP distortion in both simulated and real data containing the step-drift artifact. Together, these findings provide practical recommendations for both preventing and correcting step-drift artifacts and underscore the need for formal definition and routine recognition of this artifact to improve reproducibility and data quality in TMS-EEG research.

## Introduction

Transcranial magnetic stimulation (TMS) is a non-invasive brain stimulation method which delivers a brief magnetic pulse over the scalp, resulting in an electric field in the underlying cortex via electromagnetic induction (Barker et al., 1985). At sufficient intensities, TMS causes depolarisation of cortical neurons underneath the stimulating coil (Rothwell et al., 1991) and electroencephalography (EEG) can capture the resulting cascade of neural activity which is measured as a series of TMS-evoked potentials (TEPs) (Ilmoniemi et al., 1997). TEPs consist of positive and negative deflections at distinct latencies and polarities, with the amplitude and latency of each peak thought to reflect underlying neurophysiological processes both in the stimulated region and in broader neural networks via transsynaptic neurotransmission (Bergmann et al., 2016; Farzan et al., 2016; Rogasch and Fitzgerald, 2013; Siebner et al., 2009). In addition, TEPs also include sensory event-related potentials (ERPs) caused by the clicking noise and sensation accompanying a TMS pulse (Biabani et al., 2024, 2019; Conde et al., 2019; Gordon et al., 2021; Nikouline et al., 1999; Rocchi et al., 2021). TEPs have shown enormous potential both as a research tool to investigate healthy brain function and as a clinically relevant biomarker for diagnosing brain disorders, assessing pharmacological effects on the brain, and monitoring treatment responses (Tremblay et al., 2019; Ziemann et al., 2026).

Despite the growing interest in TEPs, TMS-EEG research is hampered by the presence of numerous different artifacts which obscure the neural signal of interest (Hernandez-Pavon et al., 2022). Several sources of artifact arise from interactions between the TMS pulse and the EEG recording system during data acquisition, including: 1) a TMS pulse artifact caused by the electric current passing through the TMS coil (Veniero et al., 2009); 2) a discharge/decay artifact caused by the dissipation of capacitances stored at the gel-skin-electrode interface (Freche et al., 2018; Julkunen et al., 2008); and 3) a recharge artifact caused when the TMS capacitors recharge between pulses (Rogasch et al., 2013; Veniero et al., 2009). Furthermore, additional artifacts arise following interaction between the TMS pulse and off-target biological systems including: 1) TMS-evoked muscle artifacts resulting from direct or indirect stimulation of craniofacial muscles by the TMS pulse (Hernandez-Pavon et al., 2012; Mutanen et al., 2013; Rogasch et al., 2013); 2) blinks and eye-movements time-locked to the TMS pulse (Bruckmann et al., 2012; Rogasch et al., 2014); and 3) neural activity in sensory and salience networks caused by auditory and somatosensory input accompanying the TMS pulse (Biabani et al., 2024, 2019; Conde et al., 2019; Gordon et al., 2021; Nikouline et al., 1999; Rocchi et al., 2021). Together, these artifacts are well recognised within the TMS-EEG literature and have guided the development of targeted online (i.e., during data acquisition) and offline (i.e., during data processing) suppression methods such as dedicated acquisition methods and cleaning pipelines for TMS-EEG data (Cline et al., 2021; Hernandez-Pavon et al., 2023, 2022; Ilmoniemi et al., 2015; Rogasch et al., 2017). However, continued refinement of artifact characterisation and suppression remain a central methodological priority to further improve TMS-EEG data quality.

One source of TMS-EEG artifact which has received relatively less interest is persistent offsets and drifts in EEG voltage which can follow the TMS pulse. This ‘step-drift’ artifact presents as an offset of up to 10 µV in amplitude following the TMS pulse which can last several seconds before slowly drifting towards pre-stimulus amplitude levels and appears separate from discharge artifacts (Rogasch et al., 2013). Throughout the manuscript, we refer to this artifact interchangeably as either the step or step-drift artifact. The source of the step-drift artifact remains unclear. One possibility is that the artifact may represent an interaction with online filters. Most TMS-compatible EEG systems operate with either AC-or DC-coupled inputs prior to digitisation. AC coupling typically applies a high-pass filter in the range of ∼0.016–0.1 Hz to suppress slow drifts and maintain the signal within the optimal dynamic range of the amplifier, but this comes with the risk of introducing additional spurious trends/drifts in the signal. By comparison, DC coupling does not require a high-pass filter which preserves slow signal components but carries an increased risk of baseline drift and amplifier clipping (Hernandez-Pavon et al., 2023). Recording using DC-coupling without high-pass filtering is typically recommended for TMS-EEG data collection (Hernandez-Pavon et al., 2023), however some studies use a very low high-pass filter (e.g., 0.016 Hz) with AC-coupling (Bonato et al., 2006; Veniero et al., 2009). Alternatively, the TMS pulse shape may also contribute. In DC-coupled EEG recordings made from a melon to isolate non-biological artifacts, Rogasch and colleagues reported anecdotal evidence that monophasic pulses from several different Magstim and MagVenture devices resulted in a persistent offset which was less evident when using biphasic pulses (Rogasch et al., 2013). However, systematic studies comparing the effect of different amplifier settings and pulse shapes on step artifacts in humans have not been reported.

The presence of step-drift artifacts in TMS-EEG signals introduces additional artifacts during data processing. For example, distinct electrical shapes within the EEG signal can yield characteristic artifacts when processed through common filters. Sharp, step-like deflections and slow drift components contain broad spectral content, meaning that when they pass through band-pass filters they generate predictable distortions such as ringing, edge effects, and spurious oscillatory activity (De Cheveigné and Nelken, 2019; Widmann et al., 2015). These distortions arise not from neural activity but from the interaction between the non-sinusoidal shape of the artifact and the filter’s impulse response (Tanner et al., 2016, 2015), highlighting the importance of understanding artifact morphology when interpreting filtered TMS-EEG data. Importantly, processing pipelines with specific steps which differ from standard TMS-EEG pipelines may be required to prevent additional filter-related artifacts if step-drift artifacts are present in TMS-EEG data (de Cheveigné and Arzounian, 2018).

In this study, we report data collected by our group in which step-drift artifacts were unexpectedly identified, from which three primary aims were derived. First, we describe the presence of step-drift artifacts in TMS-EEG recordings and use simulated data to characterise additional filtering artifacts introduced by steps-drifts in TMS-EEG signals. Second, we investigate the impact of different methodological choices during data collection on the size of step-drift artifacts, including different pulse shapes (monophasic vs. biphasic) and online high-pass filter settings (AC-coupled vs. DC-coupled). Third, we assess different combinations of data processing methods to determine the optimal steps for mitigating filtering artifacts when TMS-EEG data contain step artifacts. In the first stage of this analysis, we use simulated data with a known ground truth to evaluate the efficacy of different temporal filter and detrending combinations on minimising filtering artifacts. We then compare the optimal processing pipeline we identified to another common pipeline using real TMS-EEG data with step artifacts to assess the impact of pipeline choice on TEPs.

## Methods

### Participants

Sixteen healthy adults were recruited via university email lists and flyers to participate in an unrelated study investigating the effect of coil positioning and auditory masking methods on TEPs (experiment 1). One participant withdrew due to discomfort associated with auditory masking, yielding a final sample of fifteen participants (11 females; mean age = 30.3 ± 7.6 years, range = 23-47 years). After noticing the step artifact in the data set and determining an incorrect amplifier configuration was used for data collection (see *experiment 1* details), four participants returned to participate in an additional experiment to assess the impact of amplifier settings and pulse shape on step artifacts (*experiment 2*), and two participants completed another experiment assessing the impact of amplifier range settings on clipping artifacts (*experiment 3*). All participants provided written informed consent to procedures approved by The University of Adelaide Human Research Ethics Committee (H-2019-220) and experiments were performed in accordance with the National Statement on Ethical Conduct in Human Research from the National Health and Medical Research Council (2018). Inclusion criteria required no history of neurological or psychiatric disorders and no current psychiatric medications. Standard safety screening was conducted to exclude any contraindications to MRI or TMS (e.g., metal implants, history of seizures).

### Experimental procedures

*MRI:* All participants underwent a T1-weighted MRI scan prior to TMS-EEG sessions for use in neuronavigation of the TMS coil (3T Siemens Magnetom Cima.X; MPRAGE, TA: 5.12 min, TR: 2300 ms, TE: 2.9 ms, TI: 900 ms, voxel size: 1 mm^3^, flip angle: 8, slices: 192). T1-weighted images underwent cortical surface reconstruction using FreeSurfer’s *recon-all* pipeline (Fischl, 2012). Brodmann area 8 was identified on each participant’s cortical surface by mapping the Brodmann atlas (Pijnenburg et al., 2021) to individual anatomy using FreeSurfer’s *mris_ca_label* function. The resulting annotations were converted to individual labels for each hemisphere and subsequently transformed into a volume in native space using *mri_label2vol* for display in the neuronavigation software.

### Experiment 1 - Prefrontal TMS-EEG with Optimised Coil Positioning and Auditory Masking

Participants completed 6 different conditions designed to assess variations in coil positioning and auditory masking methods on TEPs following stimulation of prefrontal cortex, the details and results of which will be reported elsewhere. For the current study, data from 1 of these conditions were analysed. Transcranial magnetic stimulation was delivered using a figure-of-eight coil connected to a Magstim200 stimulator. Resting motor threshold (RMT) was determined individually for each participant by stimulating the left primary motor cortex and identifying the minimum intensity that elicited motor-evoked potentials (MEPs) greater than 0.05 mV in the right first dorsal interosseous (FDI) muscle recorded via surface electromyography, in at least 5 out of 10 consecutive trials (Rossini et al., 1994). Coil orientation during motor hotspotting was maintained at approximately 45° relative to the mid-sagittal plane. Pulses were monophasic with current direction running posterior-anterior (PA) in the underlying cortex. For TMS-EEG data collection, the TMS coil position was set to target Brodmann Area 8 in the left hemisphere corresponding to superior frontal gyrus. Neuronavigation was used to track coil position in relation to the underlying cortex (BrainSight^TM^, Rogue Resolutions). Coil position and angle were further optimised to maximise the amplitude of the P20/N40 TEP and minimise TMS-evoked muscle artifact using a customised optimisation process which included online feedback collected with the rt-TEP toolbox (Casarotto et al., 2022) and custom MATLAB scripts. A grid of four stimulation sites separated by ∼10 mm was placed within BA8 and 15 pulses was collected at different coil angles (60°, 90°, 120° to the midline) from each site. TMS intensity was set to 120% of each participant’s RMT adjusted for coil-to-cortex distance at each individual site (Stokes et al., 2005). The site/angle with no/ minimal muscle artifact and the largest peak-to-peak P20/N40 TEP amplitude was selected as the site for stimulation.

EEG was recorded using a 64-channel cap (multitrode electrode for TMS, EasyCap GmbH) connected to a TMS-compatible amplifier (BrainAmp MR Plus, Brain Products GmbH). Electrodes were positioned according to the international 10–20 system. A ground electrode was placed on the forehead, with a reference electrode positioned adjacent to it. Impedances for all electrodes were kept below 10 kΩ prior to recording and stimulation. EEG data were sampled at 5,000 Hz. Although our group typically use DC-coupling for TMS-EEG experiments (Biabani et al., 2024, 2019), we mistakenly used a configuration file with default amplifier settings including AC-coupling (online filters = 0.016-1000 Hz; range = ± 3.28 mV). 120 TMS pulses were delivered (ISI = 5 s; 4-6 s jitter) with participants sitting at rest with eyes open during data collection.

During recording, masking noise was continuously played to participants using in-ear earphones covered by ear defenders to minimise perception of the TMS click sound. A customised auditory masking sound was generated for each participant using the TAAC toolbox prior to data collection (Russo et al., 2022). The relevant contribution of white noise and click-based noise was varied until the optimal combination to minimise click perception was identified for each participant and used throughout data collection.

### Experiment 2-Effects of Pulse Waveform and Amplifier Coupling on TMS-EEG Artifacts

The experimental arrangement was identical to experiment 1. Participants completed 4 conditions to test the impact of pulse type (monophasic vs biphasic) and amplifier settings (AC-coupling with high-pass filter [0.016-1000 Hz] vs DC-coupling with no high-pass filter [DC-1000 Hz]) on the presence of step artifacts. Combinations included: 1) monophasic pulses + AC-coupling; monophasic pulses + DC-coupling; biphasic pulses + AC-coupling; biphasic pulses + DC-coupling. For the biphasic condition, the TMS coil was connected to a Magstim Rapid^2^ using a connector. Current direction was PA-AP in the underlying cortex.

### Experiment 3 - Impact of Amplifier Dynamic Range on Clipping in DC-Coupled TMS-EEG

The experimental arrangement was again identical to experiment 1, except DC-coupling with no high-pass filter was used for the amplifier setting. Participants completed 2 conditions to test the impact of amplifier range settings on clipping artifacts when using DC-coupling. One condition used the standard amplifier range (± 3.28 mV) while a second condition used an extended range (± 16.38 mV), acknowledging the trade-off that increasing the dynamic range reduces the effective digitization resolution and therefore may diminish signal fidelity for low-amplitude fluctuations.

### Artifact examples

First, we provided an example of step, drift and filtering (i.e., ringing) artifacts observed in an individual participant from experiment 1. Data were analysed using EEGLAB (Delorme and Makeig, 2004) and TESA (v1.1.1) (Rogasch et al., 2017) in MATLAB (MathWorks Inc.) using a minimal processing pipeline (supplementary methods pipeline 1). Both the global mean field amplitude (GMFA; the standard deviation of EEG amplitude across channels at each timepoint) and a butterfly plot showing all channels were plotted. To help demonstrate the temporal and spatial characteristics of the step-drift artifact, FastICA (Hyvärinen and Oja, 2000) was run on the data and the independent component representing the step-drift artifact (IC1) was selected and plotted. We then applied a band-pass Butterworth filter (1-80 Hz, second order) to the data to demonstrate additional ringing and edge artifacts introduced by filtering over the step and drift artifacts.

To further demonstrate the interaction between the different sources of artifact and offline filters, we simulated a step artifact by generating a flat line (amplitude = 0 µV) equivalent in length to our EEG epochs (-1000 to 1000 ms) and then added a constant value (amplitude = 5 µV) to the post-stimulus window (0 to 1000 ms). We then separately applied a high-pass (1 Hz, second order) and low-pass (80 Hz, second order) Butterworth filter to the simulated data and plotted the outcomes. We repeated the analysis on data with a simulated drift artifact to compare outcomes (linear drift rate =-5 µV/s; starting amplitude = 5 µV).

### Testing experimental arrangements to minimise step artifacts (online)

To assess the impact of online methods (AC-vs. DC-coupling; monophasic vs. biphasic pulse shapes) on artifacts in data from experiment 2, we used a minimal data cleaning pipeline implemented with EEGLAB and TESA (supplementary methods pipeline 2). Step artifact amplitudes were quantified by measuring the mean of the GMFA between 50 to 500 ms (starting value chosen to avoid any large transients in the data following the TMS pulse). To assess the impact of offline filtering on the data, we also applied a second-order Butterworth band-pass filter (1-80 Hz) using the *pop_tesa_filtbutter.m* function. Ringing artifacts were quantified by measuring the mean GMFA between-100 to-2 ms (e.g., before the TMS pulse as ringing artifacts are introduced before and after step artifacts due to zero-phase filter). Due to the low sample size (n=4), the dependent variables were grouped into either AC-coupling and DC-coupling conditions, or monophasic and biphasic conditions and compared using a two-sided Wilcoxon rank sum test (α = 0.05) with two contrasts: AC-vs. DC-coupling; and mono-vs. biphasic pulse shape.

### Developing and testing cleaning pipelines to minimise filtering artifacts (offline)

#### Function development

To assess whether we could minimise filtering artifacts resulting from step artifacts during offline data cleaning we developed four new functions for the TESA toolbox: 1) *pop_tesa_robustdemean.m* which calculates the robust mean (i.e., excluding values outside a defined SD range) within a given data range and then subtracts this value from all data points within the range (e.g., fits and removes an offset from the signal). 2) *pop_tesa_robustdetrend.m* which calculates a robust best fitting polynomial of degree *n* within a given data range and then subtracts this polynomial from data points within the range (e.g., fits and removes drifts in the signal). 3) *pop_tesa_modifiedbandpassfilter.m* which was based on methods developed in the AARATEP toolbox to avoid ringing artifacts caused by high-pass filtering over large steps in the data (Cline et al., 2021). This function uses autoregressive extrapolation to artificially extend the time series forward from the start of the TMS-pulse removal window and backwards from the end of the window (e.g., by 500 ms), thereby extending and smoothing the ‘edge’ of the data. Butterworth filters are then applied separately to the extended pre-and post-data periods and the filtered data is blended back together using sigmoidal weightings over the removal window. 4) *pop_tesa_interactivechanreject.m* which generates an interactive figure for manually checking bad channels by visualising a TEP butterfly plot. Individual channels can be manually selected and removed.

#### Simulated data tests

To compare the efficacy of different combinations of detrending and filtering for removing the step artifact and preventing filtering artifacts, we generated simulated TMS-EEG data. First, we generated an ‘absolute ground truth’ TEP by adding together two damped oscillations: a beta oscillation (frequency = 25 Hz; decay constant = 0.1 s; no. cycles = 1.5; starting phase = 270 degrees; amplitude = 5 µV; onset = 10 ms) and a theta oscillation (frequency = 5 Hz; decay constant = 0.1 s; no. cycles = 1.5; starting phase = 90 degrees; amplitude = 3 µV; onset = 60 ms). The time series was then smoothed using a low-pass Butterworth filter (40 Hz, second order). The simulated TEP showed characteristics of prefrontal TEPs including peaks at P20, N45, P65, N120 and P200. We then extracted baseline data (i.e., before the TMS pulse) from the F1 channel of real TMS-EEG data collected in our experiments (n=15;-3500 ms to-500 ms; 120 trials), inserted artificial TMS triggers at the mid-point of the extracted period, epoched around the artificial triggers (-1500 to 1500 ms) and baseline corrected the data (-500 to-10 ms). Next, we added our ‘absolute ground truth’ TEP to each trial so the onset coincided with timepoint 0 (i.e., the centre of the epoch), and then removed and interpolated data around the artificial TMS trigger (-2 to 10 ms; cubic interpolation). Finally, we created five conditions with artificial step artifacts between 2 and 10 µV in amplitude (2 µV increments) by adding constant values to the post TMS data points.

We then assessed how different combinations of cleaning methods impacted our simulated TEPs with different step artifact amplitudes. Robust demeaning and detrending (polynomial order = 1) were applied separately on the pre (-1500 to-2 ms) and post (10 to 1499 ms) TMS periods. A standard high-pass Butterworth filter (1 Hz, second order), and a modified high-pass Butterworth filter (1 Hz, fourth order, autoregressive extension = 900 ms) were also used. In total, 8 combinations of methods were tested including:

1. robust demean;
2. robust detrend;
3. standard Butterworth filter;
4. modified Butterworth filter with autoregressive extrapolation;
5. robust demean + Butterworth filter;
6. robust detrend + Butterworth filter;
7. robust demean + modified Butterworth filter;
8. robust detrend + modified Butterworth filter.

We first quantified the impact of increasing step artifact amplitude on TEP distortion by calculating the root mean square error (RMSE; from 10 to 300 ms) between simulated TEPs with no step artifact following cleaning compared to those with increasing step amplitudes. Next, we compared how well each method retrieved the ‘absolute ground truth’ TEP by calculating the RMSE between each cleaning combination and the ground truth (between 10 to 50 ms). Cleaning combinations were then ranked from highest to lowest RMSE. The approach was repeated for the 0 and 10 µV step artifact conditions and the cleaning combination with the lowest RMSE in the 10 µV step artifact condition chosen as the optimal pipeline. To assess how these cleaning approaches performed in the presence of other non-linear artifacts common in TMS-EEG data, we repeated the above analyses introducing a decay/discharge artifact at timepoint 0 for the 0 and 10 µV step artifact conditions. Decay/discharge artifacts were simulated as decay following a second-order power law (duration = 500 ms; scaling coefficient = 0.005 µV.s^2^; exponent = 2).

#### Real data tests

Having established the optimal cleaning pipeline for minimising step and ringing artifacts in simulated data, we then compared this modified pipeline to a standard cleaning pipeline in real data from experiments 1 and 2. The standard and modified pipelines are summarised in table 1. The standard pipeline consisted of the following. First, we removed and interpolated data around the TMS pulse in the continuous recordings (-2 to 10 ms; cubic interpolation) and resampled the data (1000 Hz). Channels showing flat lines longer than 4 s were removed, data were epoched around the TMS pulse (-1000 to 1000 ms) and baseline corrected (-500 to-10 ms), and epochs with values >3 SD either across a single channel or across all channels were removed. Any missing channels were then replaced using spherical interpolation and the data were referenced to the average of all channels. Butterfly plots of TEPs were manually checked at this point and bad channels were identified and removed using *pop_tesa_interactivechanreject.m*. Next, interpolated data and channels were removed prior to FastICA and independent components representing TMS-evoked muscle activity were automatically detected and removed using the default values of the *pop_tesa_compselect.m* function. Removed data and channels were interpolated and Butterworth band-pass (1-80 Hz, second order) and band-stop (48-52 Hz, second order) filters were applied. Interpolated data and channels were again removed prior to a second round of FastICA and independent components representing residual TMS-evoked muscle activity, blinks, eye movement and ongoing muscle activity were automatically detected and removed using the default values of the *pop_tesa_compselect.m* function. Finally, removed data were interpolated and the data were again referenced to the average of all channels. The modified pipeline was identical to the standard pipeline except the band-pass and notch filtering steps were replaced by: 1) a robust detrend (degree = 1; equivalent to a linear detrend) performed separately on the pre (-1000 to-2 ms) and post (10 to 999 ms) time windows; 2) cubic interpolation of the TMS-pulse removal window; 3) a modified high-pass Butterworth filter with autoregressive extrapolation (1 Hz, fourth order, extension = 900 ms); 4) a standard low-pass filter (80 Hz, second order); and 5) a modified notch filter (48-52 Hz, fourth order, extension = 900 ms). We chose a modified notch filter as we observed additional ringing artifacts associated with the standard Butterworth filter in testing (supplementary figure S1).

**Table 1:** summary of the standard and modified processing pipelines. The modified pipeline was identical to the standard pipeline except at steps 15a-e.

| Step | Standard pipeline | Modified pipeline |
| --- | --- | --- |
| 1 | Remove TMS pulse (-2 to 10 ms). |  |
| 2 | Interpolate missing data (cubic). |  |
| 3 | Downsample (1,000 Hz). |  |
| 4 | Remove channels with flat lines. |  |
| 5 | Epoch (-1000 to 1000 ms). |  |
| 6 | Baseline correct (-500 to -10 ms). |  |
| 7 | Remove epochs >3 SD in amplitude |  |
| 8 | Replace missing channels (spherical interpolation). |  |
| 9 | Re-reference to average of all channels. |  |
| 10 | Manually check removed channels. |  |
| 11 | Remove interpolated data. |  |
| 12 | FastICA. |  |
| 13 | Automatically remove components representing TMS-evoked muscle activity. |  |
| 14 | Interpolate missing data. |  |
| 15a | Butterworth band-pass filter (1-80 Hz, second order) | Robust detrend (degree = 1) on pre and post TMS periods separately. |
| 15b | Butterworth band-stop filter (48-52 Hz, second order). | Interpolate missing data (cubic). |
| 15c |  | Modified high-pass Butterworth filter with autoregressive extrapolation (1 Hz, fourth order, extension = 900 ms). |
| 15d |  | Standard low-pass filter (80 Hz, second order) |
| 15e |  | Modified Butterworth band-stop filter (48-52 Hz, fourth order, extension = 900 ms). |
| 16 | Remove interpolated data. |  |
| 17 | FastICA. |  |
| 18 | Automatically remove components representing TMS-evoked muscle, eye blinks, eye movement, ongoing muscle activity. |  |
| 19 | Interpolate missing data |  |
| 20 | Re-reference to average. |  |

To assess the impact of cleaning pipelines on ringing artifacts in data with and without step artifacts (experiment 2), we calculated the GMFA in the baseline period before the TMS pulse (-100 to-2 ms) to quantify the ringing artifact in the AC-(step present) vs DC-coupling (step absent) dataset (n=4). We then grouped and compared ringing artifact amplitudes across four separate contrasts using two-sided Wilcoxon rank sum tests (α = 0.05): data with step vs. no-step using the standard pipeline; data with step vs. no-step using the modified pipeline; standard vs. modified pipeline in data with step artifacts; and standard vs. modified pipeline in data without step artifacts. To assess the impact of step artifacts and cleaning pipelines on TEP spatiotemporal characteristics, we compared TEP amplitudes in the original dataset (n=15; AC-coupling with step artifacts; experiment 1) cleaned with the standard vs. modified pipeline using cluster-based permutation statistics in FieldTrip (Oostenveld et al., 2011). Six separate time windows were assessed covering canonical TEP peaks and EEG amplitudes were average across time within each window: baseline (-110 to-90 ms); P20 (36 to 55 ms); N45 (56 to 75 ms); N120 (110 to 130 ms); and P220 (210 to 230 ms). Clusters were defined as two or more neighbouring electrodes in which the t-statistic exceeded a threshold of p < 0.05 (two-sided dependent t-test). The test statistics were calculated as the maximum of the cluster-level statistics (i.e., “maxsum”). Monte Carlo p-values were calculated on 5000 random permutations and a value of p < 0.05 (2-tailed) was used as a cluster significance threshold.

## Results

### Step, drift, and filtering artifacts

Figure 1A shows an example of step-drift artifacts in TMS-EEG data for an individual participant from experiment 1 (online settings: sampling rate = 5 kHz, AC-coupled filters, 0.016-1000 Hz). EEG amplitudes are offset immediately following the TMS pulse and do not return to baseline (average of 120 trials, epoched and baseline corrected with the pulse artifact between-2 to 10 ms removed, then re-referenced to average. F1 channel is highlighted). The step artifact offset is clearly visible in the GMFA (figure 1B). The step artifact is captured by the first independent component (figure 1C), which shows the step in the data after the pulse followed by a slow drift. The offset is largest in channels near the stimulation site (left frontal channels near F1) and can change polarity across different scalp locations. Figures 1D-E shows the TEP and GMFA following band-pass filtering (second order Butterworth filter; 1-80 Hz). Although the offset of the step artifact is mostly removed, distortions are still present immediately before and after TMS pulse (known as ‘ringing’ artifacts) and at the start and end of the epoch (known as ‘edge’ artifacts). Note that ringing artifacts occur both before and after the pulse artifact due to the zero-phase design of acausal filters which run both forwards and backwards through the data to prevent phase offsets. Edge artifacts occur due to steps in the data introduced by zero-padding at the epoch boundaries. The ringing and edge artifacts following filtering of the step-drift artifact are clearly represented in the first IC (figure 1F).

**Figure 1:**
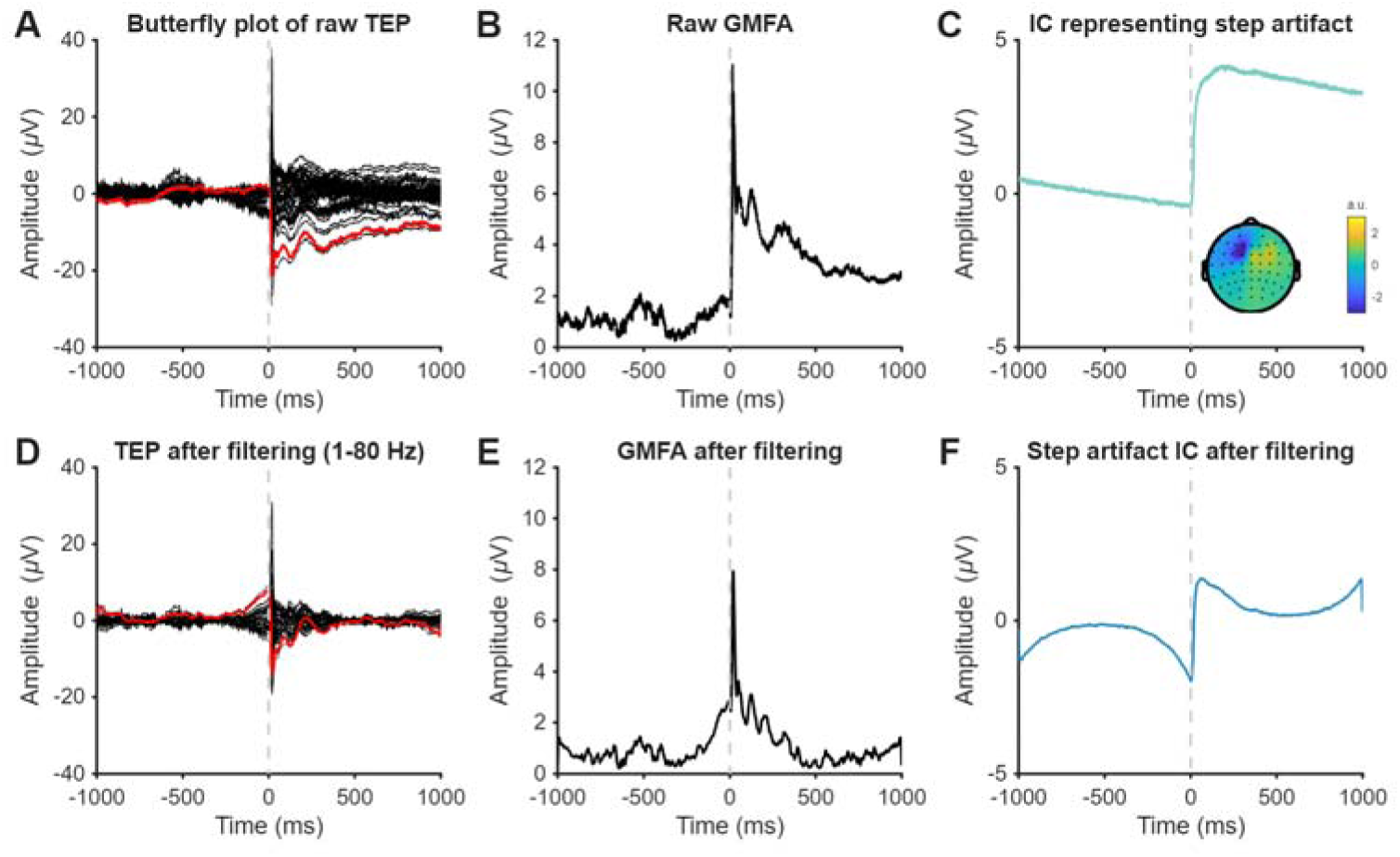
Step, drift, and filtering artifacts in TMS-EEG data. A) Butterfly plot of minimally-processed EEG data (epoched, baseline corrected, and average referenced) following a TMS pulse with a step artifact. A channel near the stimulation site (F1) is highlighted in red. Timing of the TMS pulse is indicated by grey dashed line. B) Global mean field amplitude (GMFA) of minimally-processed TEPs with a step artifact. C) Temporal (line plot) and spatial (inset topoplot) weightings of the first independent component (IC) representing the step and drift artifact. D) Same data as in A following 1-80 Hz band-pass filtering (second order Butterworth filter). E) Same data as in B following 1-80 Hz band-pass filtering (second order Butterworth filter). F) Same IC as in C following 1-80 Hz band-pass filtering.

To demonstrate the interaction between artifacts and temporal filters, we simulated a constant signal (amplitude = 0 µV) with either a step (amplitude = 5 µV; 0-1000 ms), or a drift (linear drift rate =-5 µV/s; starting amplitude = 5 µV), and then applied high-and low-pass Butterworth filters (second order) separately to the simulated data (figure 2). These analyses show that high-pass filters result in the positive/negative ringing artifacts before/after the step in the data, whereas the edge artifacts are introduced following high-pass filtering over drifts in the data. In contrast, low-pass filters have minimal impact on the data in the presence of either step or drift artifacts. Together, these analyses show that the presence of a step-drift artifact in TMS-EEG data can lead to additional filtering-related artifacts following high-pass and, by extension, band-pass filtering.

**Figure 2:**
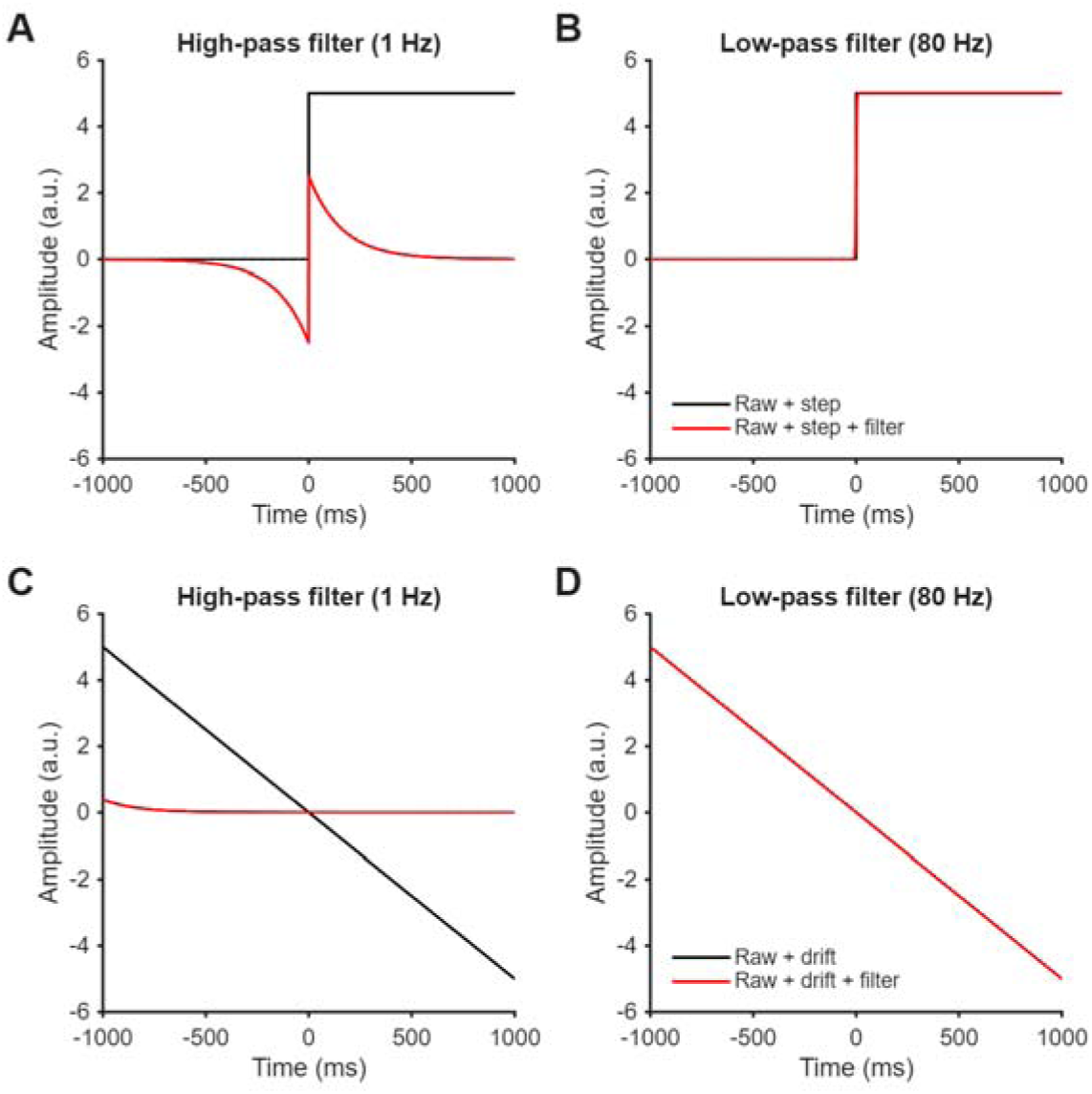
The effect of filtering over simulated steps and drifts. The impact of second order Butterworth filters on constant data with a step artifact (A, 1 Hz high-pass; B, 80 Hz low-pass) and a drift artifact (C, 1 Hz high-pass; D, 80 Hz low-pass).

### Online methods for minimising step artifacts

*Pulse shape and online filters:* Having established that the presence of step-drift artifacts can lead to additional artifacts following common filtering steps, we next assessed the impact of different online methodological choices on the amplitude of step-drift artifacts in TMS-EEG data. To assess whether the choice of online high-pass filter setting (AC-coupling with 0.016 Hz high-pass filter vs. DC-coupling with no high-pass filter) or the type of pulse shape (monophasic vs. biphasic) altered step-amplitude artifact, we compared minimally processed data from 4 participants across 4 conditions. When comparing GMFA between conditions, the post-TMS offset was larger when using AC-vs. DC-coupling (p < 0.001), but did not differ between monophasic and biphasic pulse shapes (p = 0.574; figure 3A-C). In addition, after applying a band-pass filter (1-80 Hz, Butterworth, order = 2), ringing artifacts were evident before the TMS pulse (introduced by zero-phase filter; see figure 2) and were larger with AC-coupling vs DC-coupling (p = 0.007), but did not differ between monophasic and biphasic pulses (p = 0.574; figure D-F). Together, these findings suggest that using DC-coupling with no online high-pass filters can minimise both the step artifact and subsequent ringing artifacts introduced by offline high-pass filters compared to AC-coupling with online high-pass filtering.

**Figure 3:**
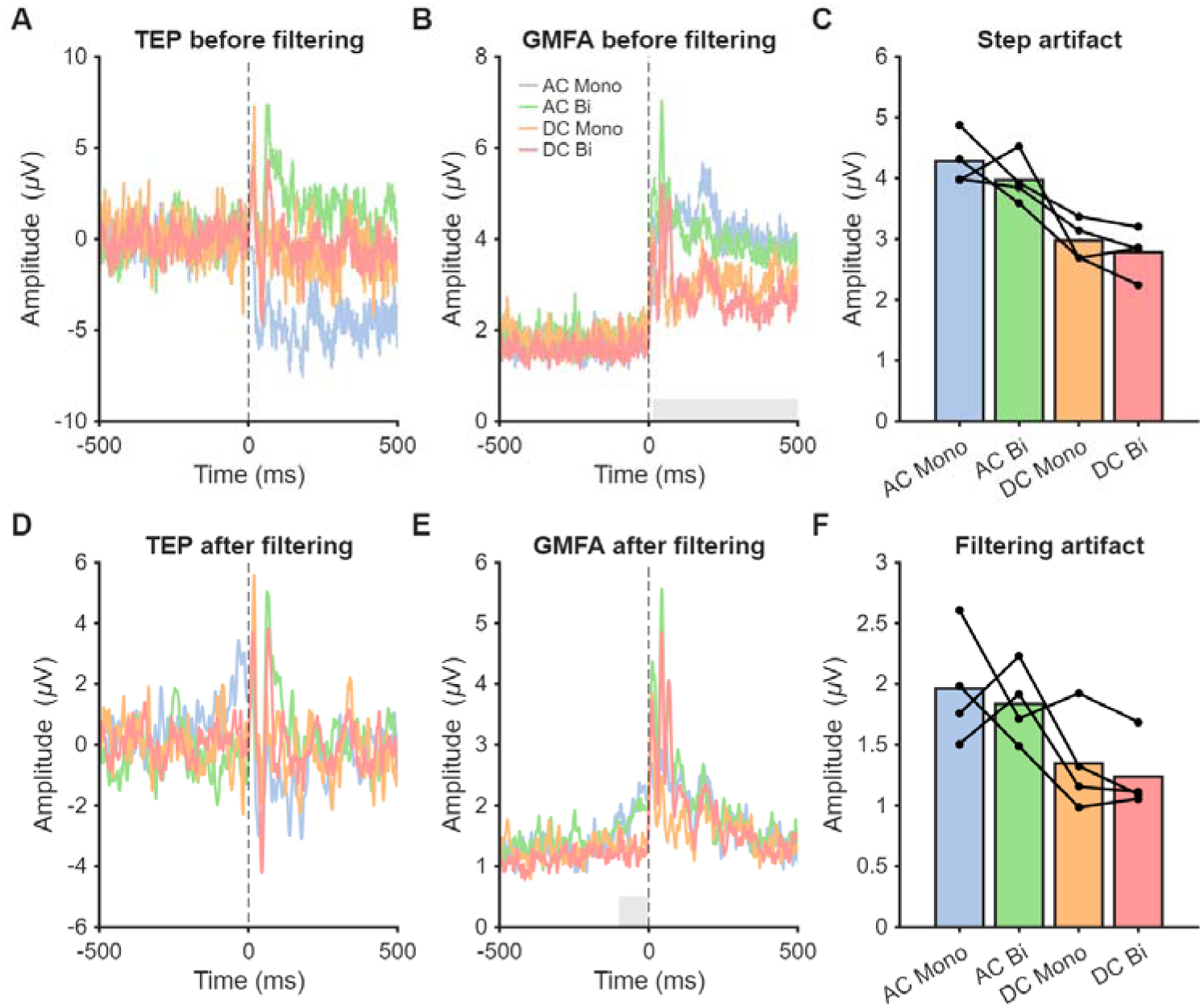
The effect of amplifier settings and TMS pulse shape on step and ringing artifacts. A) TMS-evoked potentials (TEPs) from the F3 channel averaged across participants for each condition before filtering. B) Global mean field amplitude (GMFA) averaged across participants for each condition before filtering. The grey bar represents the analysis period for quantifying the step artifact (50 to 500 ms). C) Step artifacts across different experimental arrangements. Lines represent values from individual participants, bars represent the group mean. D) TEPs averaged across participants for each condition after band-pass Butterworth filtering (1-80 Hz, second order). E) GMFA averaged across participants for each condition after band-pass Butterworth filtering (1-80 Hz, second order). The grey bar represents the analysis period for quantifying the filtering artifact (-100 to-2 ms). F) Ringing artifacts across different experimental conditions. Mono = monophasic TMS pulses; bi = biphasic TMS pulses; AC = alternating current-coupled amplifier (high-pass filter = 0.016 Hz); DC = direct current-coupled amplifier (no high-pass filter).

### Clipping

During experiment 2, we noticed a number of channels ‘clipped’ (e.g., moved outside of the operating range) (figure 4A). Instances of clipping were considerably higher in conditions using DC-coupling, particularly with monophasic pulses (figure 4B), and instances tended to increase over the duration of the recording condition (supplementary figure S2). Increasing the amplifier range from ± 3.28 mV to ± 16.38 mV reduced the instances of clipping during DC-coupled recordings for both monophasic and biphasic pulses (figure 4C), with only one channel exceeding the amplifier range across the two participants and four conditions tested (supplementary figure S3). In summary, these findings suggest caution is required using DC-coupling due to the increased risk of channels drifting outside of the amplifier recording range. However, using a larger amplifier range when collecting TMS-EEG data with DC-coupling can mitigate data lost to clipping.

**Figure 4:**
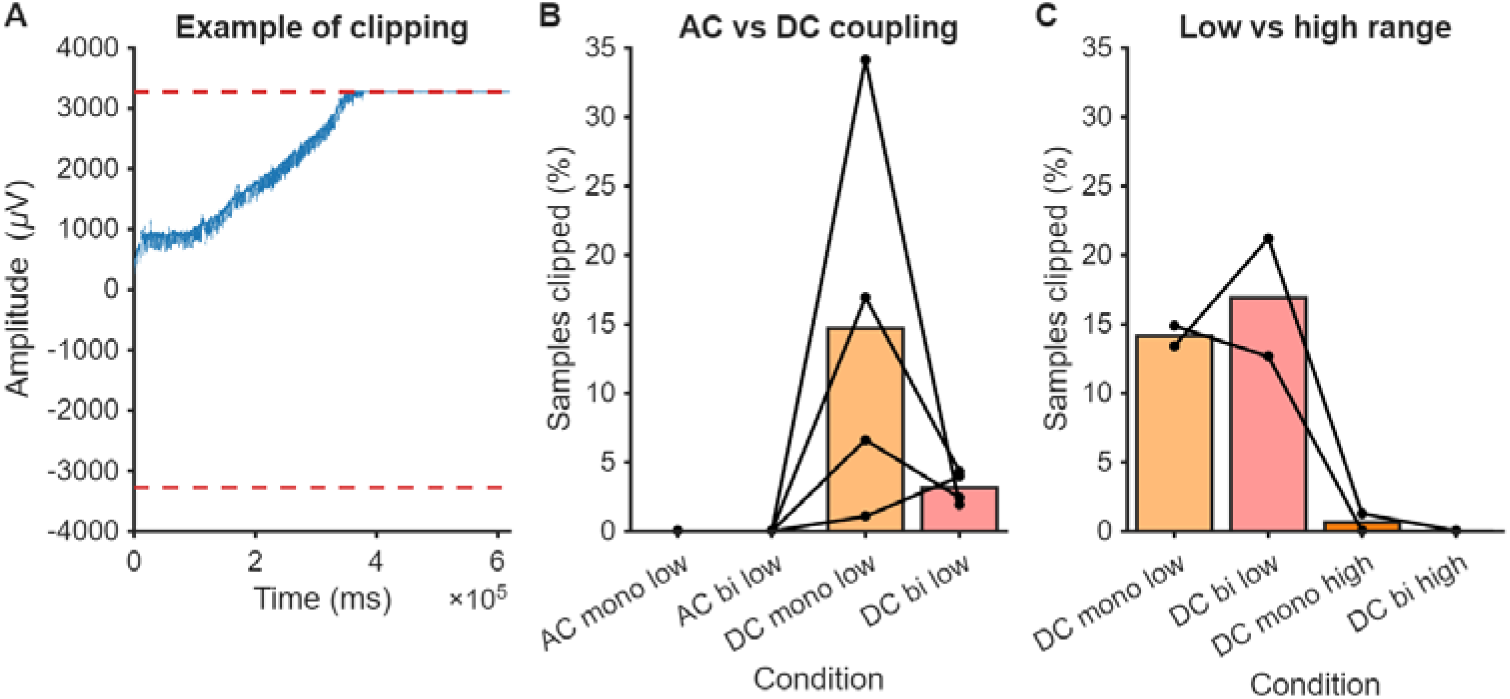
‘Clipping’ artifacts in TMS-EEG recordings. A) An example of an EEG channel that drifts outside of the amplifier recording range indicated by red dashed lines (i.e. ‘clips). The recording was made using DC-coupling (i.e., no online high-pass filter). Note that TMS pulses were removed and interpolated for visualisation. B) Percentage of samples outside of the amplifier recording range across all channels for each condition. Lines represent values from individual participants, bars represent the group mean. Mono = monophasic TMS pulses; bi = biphasic TMS pulses; AC = alternating current-coupled amplifier (high-pass filter = 0.016 Hz); DC = direct current-coupled amplifier (no high-pass filter). C) Percentage of samples outside the recording range. Lines represent values from individual participants (high = ± 16.38 mV low = ± 3.28 mV).

### Offline methods for minimising filtering artifacts

#### Simulations

To assess the impact of high-pass filtering on TEPs with step artifacts, we applied a 1 Hz high-pass filter (second order Butterworth) to a simulated TEP mixed with real EEG data (n=15) and introduced simulated step artifacts of varying amplitude (figure 5A-B). Compared to no step artifact, increasing step amplitude resulted in increasing ringing artifact distortions in both the pre-stimulus and post-stimulus windows following a standard Butterworth filter (figure 5B-C). We then assessed whether using either robust demeaning or robust detrending to the pre and post TMS time window separately prior to filtering improved filter outcome. Using either method removed the distortion with increasing step amplitude (figure 5C-D). We also assessed whether using a modified Butterworth filter which uses autoregression to artificially extend the pre and post TMS windows prior to filtering improved filter outcomes. The distortion with increasing step amplitude was reduced using the modified Butterworth filter relative to a standard Butterworth filter, but was optimal when either robust demeaning or detrending were applied as well (figure 5E-F).

**Figure 5.**
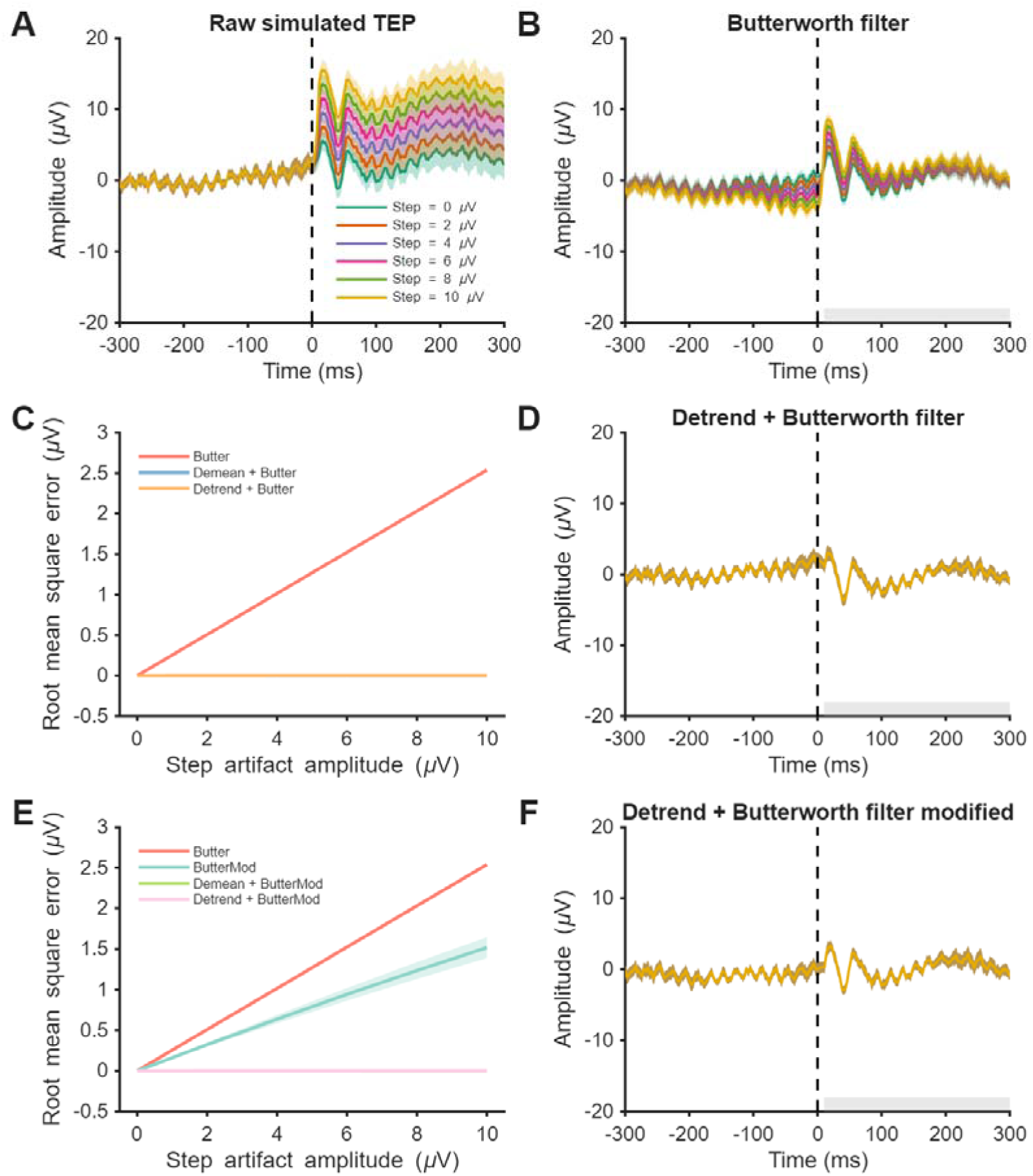
The impact of different pre-processing methods on ringing artifacts with increasing step artifact amplitude. A) Simulated TEPs embedded in real EEG data with differing simulated step artifact amplitudes. Solid line indicates group mean and shaded lines indicate standard error (n=15). Simulated TEPs following high-pass Butterworth filter (1 Hz, second order). Ringing artifacts are evident before and after the simulated TMS pulse and increase with increasing step artifact amplitude. The grey bar represents the time period used to compare root mean square error between conditions. The impact of including either robust demeaning or robust detrending prior to Butterworth filter on ringing artifacts with increasing step artifact amplitude. TEP distortion is quantified by calculating the root-mean square error with the filtered TEP with no step artifact. Note that Demean + Butter and Detrend + Butter are overlapping. D) Simulated TEPs following robust detrending and a high-pass Butterworth filter. Ringing artifacts are no longer evident with increasing step artifact amplitude. E) The impact of using a modified Butterworth filter with autoregressive extrapolation on ringing artifacts with increasing step artifact amplitude. Note that Demean + Butter and Detrend + Butter are overlapping. F) Simulated TEPs following robust detrending and a modified high-pass Butterworth filter.

To determine which combination of pre-processing methods best recovered the simulated TEP, we calculated RMSE relative to the absolute ground truth TEP (10 to 50 ms) following 8 different combinations of filters and detrending methods for both no step artifact and a 10 µV step artifact. We then sorted methods based on largest to smallest RMSE. For both cases, pre-processing generally reduced RMSE relative to raw data. However, the combination of robust demeaning/detrending and the modified Butterworth filter resulted in the smallest RMSE both with and without a step artifact (figure 6), indicating this combination is optimal for reducing drifts in the signal regardless of the presence of steps.

**Figure 6:**
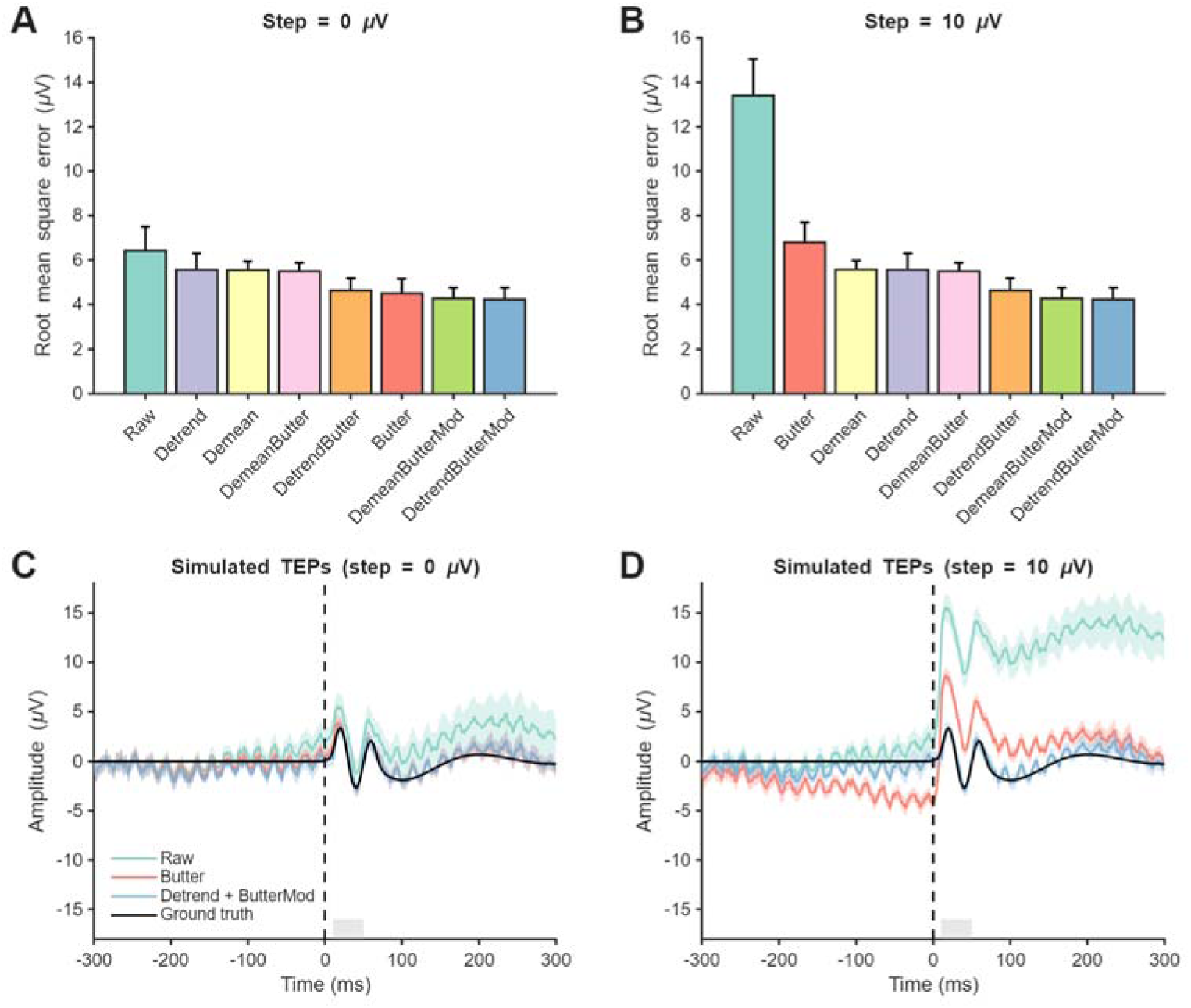
The impact of different cleaning combinations on recovering the ground truth TEP. A) Root mean square error (RMSE) between the ground truth TEP and cleaned simulated TEPs embedded in real EEG data with no step artifact for 8 different combinations of detrending and filtering. Bars represent the group mean and error bars the standard error of the mean (n=15). Cleaning combinations are ranked from highest to lowest RMSE. B) RMSE between the ground truth TEP and cleaned simulated TEPs embedded in real EEG data with 10 µV step artifact for 8 different combinations of detrending and filtering. C) Comparison of simulated TEPs embedded in real EEG data with no step artifact including raw data, data following Butterworth filtering, and the optimal cleaning combination (detrend + modified filter) vs. ground truth TEP. Solid lines represent group mean and shaded bars represent standard error. The grey bar indicates the time period in which RMSE was calculated. D) Comparison of simulated TEPs embedded in real EEG data with 10 µV step artifact including raw data, data following Butterworth filtering, and the optimal cleaning combination (detrend + modified filter) vs. ground truth TEP.

To assess how robust detrending and the modified filter performed in the presence of non-linear artifacts common in TMS-EEG recordings, we repeated the above analysis on simulated data including a discharge/decay artifact introduced at timepoint 0. The differences in RMSE between standard Butterworth filters and the optimal cleaning combination (detrend + modified filter) were similar with a discharge/decay artifact to without (supplementary figure S4), suggesting this approach is unaffected by non-linear artifacts. In summary, the combination of robust detrending and a modified band-pass filter with autoregressive interpolation appears optimal for reducing filtering artifacts in TMS-EEG data containing step artifacts regardless of the presence of other artifacts.

#### Comparison of cleaning pipelines

The results of the simulation analysis suggested that a combination of robust detrending and a modified Butterworth filter were optimal to minimise offline filtering artifacts if step artifacts were present in TMS-EEG data. To test this, we assessed whether a cleaning pipeline using the modified detrend/filtering approach was able to minimise artifacts compared with a standard cleaning pipeline in real TMS-EEG data (AC/DC-coupling data; experiment 2). As in the earlier analysis, ringing artifacts were larger in data collected with AC-coupling (i.e., with a step artifact) vs. DC-coupling (i.e., without a step artifact) using the standard pipeline (p = 0.003), but did not differ between conditions when using the modified pipeline (p = 0.645; figure 7). When directly comparing the pipelines in the data collected with AC-coupling, the modified pipeline reduced the amplitude of the ringing artifact compared with the standard pipeline (p = 0.005). In contrast, there was no difference when comparing pipelines in data collected with DC-coupling (p = 0.505). In summary, these analyses suggest that the modified cleaning pipeline is able to both suppress the step artifact and reduce subsequent ringing artifacts caused by offline filtering, resulting in TEPs that are similar to those collected without a step artifact.

**Figure 7:**
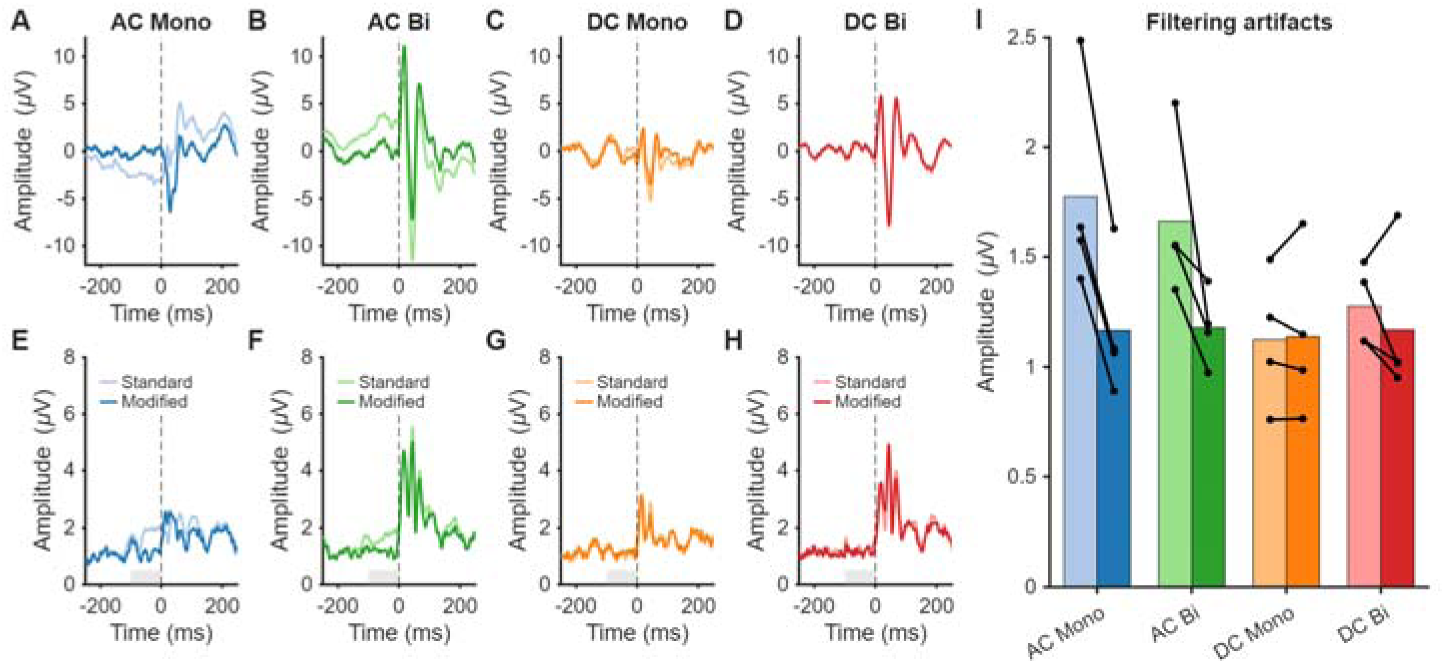
Comparison of a standard and modified cleaning pipeline for minimising ringing artifacts in data with and without step artifacts. A-D) TMS-evoked potential (TEP) amplitude from data following standard and modified cleaning pipelines across different experimental settings. Solid lines represent group means (n=4). E-H) Global mean field amplitude (GMFA) from data following standard and modified cleaning pipelines across different experimental settings. The grey bar represents the analysis period for quantifying the filtering artifact (-100 to-2 ms). I) Ringing artifact amplitude from GMFA data following standard and modified cleaning pipelines across different experimental settings. Bars represent group means and lines represent values from individual participants.

Finally, we assessed the generalisability of this approach by comparing the standard vs. modified cleaning pipeline in a larger group of participants using a sub-optimal recording arrangement resulting in step artifacts (n=15; experiment 1). The modified cleaning pipeline resulted in substantial differences in the amplitude and spatial distribution of canonical prefrontal TEPs compared to the standard pipeline, including the P20 (p < 0.001), N45 (p < 0.001), P65 (p < 0.001), N120 (p < 0.001) and P220 (p = 0.004), as well as during baseline (p < 0.001) (figure 8). These findings demonstrate that standard Butterworth filters can introduce ringing artifacts that substantially alter the spatiotemporal properties of TEPs, especially in data with steps or offsets. Importantly, these artifacts can be minimised by using pipelines which separately detrend the pre and post data prior to filtering, and that use modified filters which take into account edges resulting from removal of the TMS pulse artifact.

**Figure 8:**
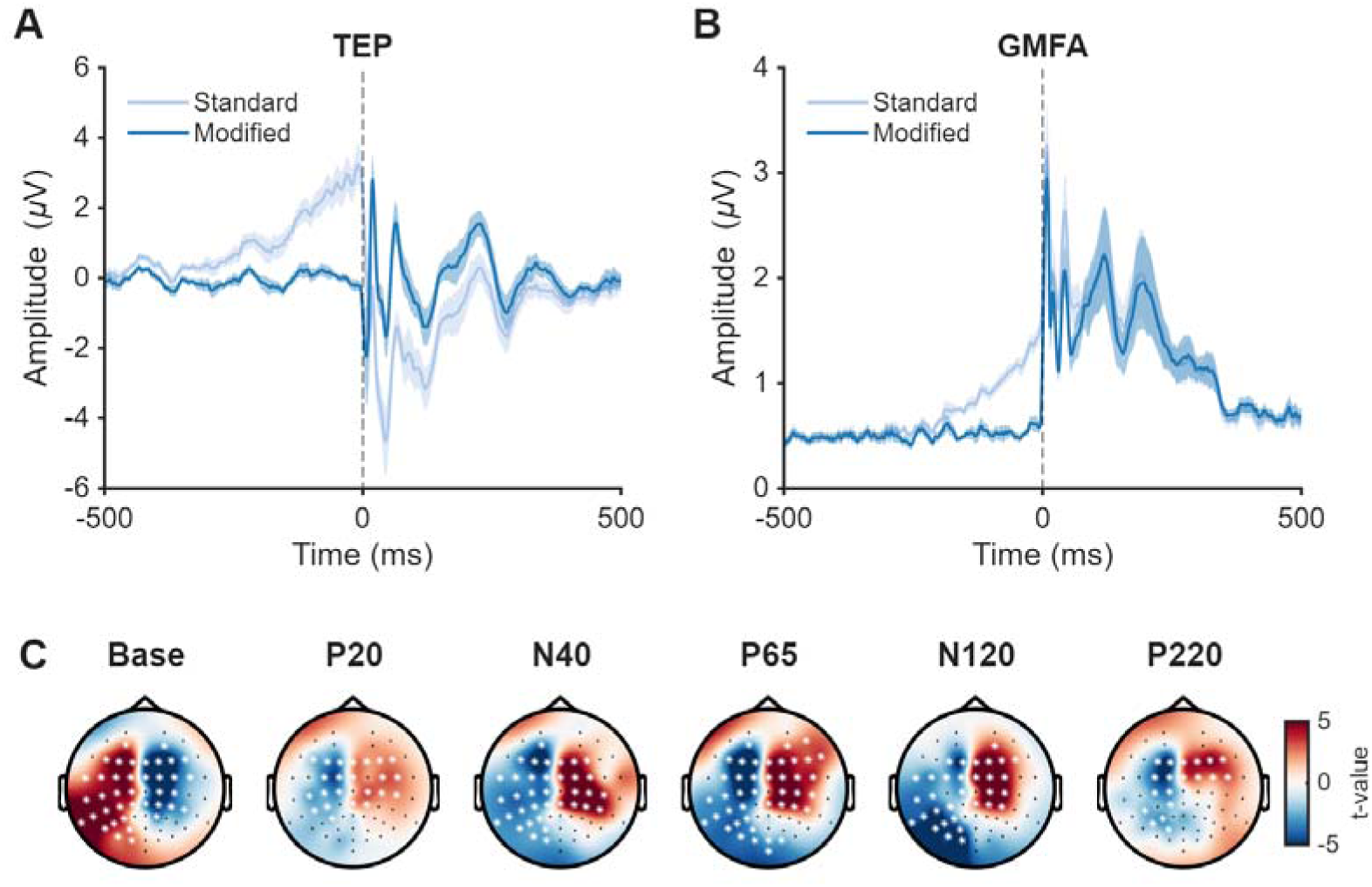
Comparison of TEPs following a standard and modified cleaning pipeline in data with a step artifact. A) TMS-evoked potential (TEP) data from a single channel near the site of stimulation (F1) following a standard and modified cleaning pipeline in data with a step artifact. Solid lines represent group mean (n = 15) and shaded bars represent standard error of the mean. B) Global mean field amplitude (GMFA) of TMS-EEG data with a step artifact following a standard and modified cleaning pipeline. C) Topoplots showing statistical maps comparing TEPs following standard vs. modified cleaning pipelines. Colours represent t-values and asterisks channels contributing to significant clusters following cluster-based permutation tests (p<0.05).

## Discussion

Artifacts are an enduring problem in TMS-EEG recordings. In the current study, we show that TMS-evoked activity can be distorted by a persistent non-neural artifact, appearing as a sudden post-TMS voltage step followed by a slow drift in EEG amplitude. Using offline filters on data containing the ‘step’ and ‘drift’ components introduce additional ringing and edge artifacts respectively, further distorting TEPs. The step-drift artifact was present exclusively under AC-coupled amplification with an online high-pass filter and absent during unfiltered DC-coupled recording, supporting a hardware–recording interaction rather than a neural origin. Such effects are therefore amenable to isolation and control under appropriate acquisition conditions. Furthermore, we identified a combination of detrending and modified filters which minimised TEP distortion in both simulated and real data containing step-drift artifacts. Collectively, these findings motivate clear recommendations for optimising online acquisition settings to mitigate the artifact and to address secondary filter-induced distortions when contamination occurs.

Previous work has characterised several artifacts directly induced by the TMS pulse in EEG recordings. Artifacts include the TMS-pulse artifact (Veniero et al., 2009; Virtanen et al., 1999), a discharge/decay artifact associated with charges stored at the electrode-gel-skin interface (Freche et al., 2018; Rogasch et al., 2013; Sekiguchi et al., 2011; Virtanen et al., 1999), and a recharge artifact associated with recharging of TMS capacitors (Rogasch et al., 2013; Veniero et al., 2009). In contrast, the present study addresses a distinct, longer lasting step-drift artifact that emerges post pulse and reflects persistent distortions in EEG voltage. Unlike discharge artifacts, which decay following a second order power law (Freche et al., 2018), recovery from the step-drift artifact is slow and non-exponential and appears as a separate component following ICA, suggesting a distinct mechanism. As step artifact amplitudes were significantly larger under AC-coupling with an online high-pass filter compared with DC-coupling with no filter (figure 3), we suggest a hardware-related origin rather than a classical discharge process, possibly reflecting an interaction between the TMS pulse and high-pass filter. Prior recommendations suggest DC-coupled systems over AC-coupled systems for TMS-EEG recordings (Hernandez-Pavon et al., 2023). Our findings provide direct evidence supporting this recommendation, suggesting that AC-coupling can induce additional TMS-related artifacts not present when using DC-coupling with no online high-pass filters.

Notably, data collected with DC coupling using the standard amplifier range (±3.28 mV) resulted in an increased number of channels where signal drifted beyond the amplifier’s dynamic range (i.e., clipped). Increasing the range from ±3.28 mV to ±16.38 mV significantly reduced channel clipping, supporting this as a more appropriate configuration. However, this improvement comes at the cost of reduced effective digitization resolution, which may attenuate the fidelity of lower-amplitude signal fluctuations. An alternative approach is to frequently reset the DC offset of the amplifier, which can be automated for certain EEG amplifiers (e.g., through the BrainVision Recorder’s COM interface). Therefore, while DC-coupling minimises step-drift artifacts, care must be taken to ensure appropriate amplification settings to avoid data lost to clipping.

Another important consideration is how TMS hardware settings such as pulse waveform shape interact with the EEG recording system to produce transient distortions. For example, one study anecdotally reported larger-amplitude step artifacts with distinct morphologies for monophasic compared to biphasic pulses (Rogasch 2013). In contrast, we observed no pulse waveform-dependent differences when amplifier settings were matched in the current study. The reasons for this discrepancy are unclear but may reflect differences in amplifier models used across studies (e.g., BrainAmp vs Compumedics). Future work directly comparing different hardware combinations are required to more formally test the generalisability of these findings.

In addition to hardware settings during data collection, digital signal processing methods applied during data cleaning can also distort the brain signal of interest. For example, it is well known that temporal filtering applied to characteristic artifact waveforms can introduce secondary, non-neural distortions, with sharp transients known to generate an artifactual oscillatory response (“ringing”) following high-pass filtering (De Cheveigné and Nelken, 2019). Depending on the characteristics of the artifact, ringing artifacts can resemble the time course of event-related potentials like TEPs (Tanner et al., 2015). Extending this principle, the step-drift morphology constitutes an additional filter-sensitive feature, whereby filtering across sustained offsets and gradual drifts yields predictable ringing and edge artifacts. This distortion differs from previously described TMS-EEG artifacts and no consensus method currently exists for its systematic removal or control.

To our knowledge, this study represents the first formal characterisation of a step-drift artifact in TMS-EEG, despite features consistent with its morphology being visible in previously published works (Rogasch et al., 2017, 2013) (TESA toolbox manual). Given its prevalence and impact, we provide practical recommendations to reduce the artifact at acquisition, alongside a targeted post processing removal strategy for cases in which it persists or inappropriate amplifier settings are used. Importantly, we validated our processing approach in both simulated and real data, in line with recent recommendations (Hernandez-Pavon et al., 2023; Rogasch et al., 2022). Ringing and edge artifacts indicate that the step component must be removed prior to filtering. Indeed, estimating and subtracting the step magnitude using a robust detrending approach, with a first-order polynomial fit applied separately to pre-and post-TMS signals, significantly reduced error relative to the ground truth in simulated data, and produced more physiologically plausible waveform shapes in real data. Error reductions scaled linearly across increasing step amplitudes, indicating consistent and proportional artifact removal. In addition to step removal, we tested a modified Butterworth filter designed to minimise ringing and edge artifacts arising from the interaction between residual offsets and temporal filtering by artificially extending the pre-and post-TMS signal using autoregressive extrapolation (Cline et al., 2021). Although improvements from the modified filter alone were modest, its inclusion further reduced filtering-related distortions when combined with robust detrending. Importantly, this optimised combination of methods was robust to other non-linear artifacts often present in TMS-EEG data, including decay/discharge artifacts. Figure 8 further demonstrates clear improvements in TEP quality when using these recommended methods on a larger data set, including reduced baseline distortions and more physiologically plausible TEP profiles compared with the standard pipeline. We also introduce an interactive channel selection tool to facilitate identification and removal of channels disproportionately affected by step-related artifacts, which we expect will yield additional improvements in data quality when applied judiciously. All methods are included in the TESA toolbox for use and additional testing by the TMS-EEG community (Mutanen et al., 2020; Rogasch et al., 2017).

A limitation of the present study is that the findings are derived from a specific set of hardware and amplifier configurations available within our laboratory. While the step-drift artifact was highly stereotyped across participants in our dataset, further validation is needed to assess its prevalence and morphology across other amplifier models. Replication across platforms will be essential to determine the generalisability of our observations and the robustness of the proposed mitigation strategies. Future work should also evaluate whether the integration of the offline correction methods proposed in our study are effective in independent data sets collected by different research groups. Incorporating our functions into the TESA toolbox provides widespread access to these methods, thereby facilitating comparisons by other groups.

Together, these findings support two overarching recommendations that prioritise prevention of step-drift distortions through optimal amplifier settings. First, record TMS-EEG data using DC-coupled amplification without high-pass filtering, and second, ensure the amplification range is sufficient to prevent clipping. We also propose a modified offline pipeline that enables targeted recovery of affected neural signals when prevention is not possible. These results underscore the importance of explicitly identifying and controlling for artifacts within TMS-EEG acquisition and processing pipelines. Accordingly, formal definition and routine recognition of step-drift artifacts are essential to improve data interpretation, reproducibility, and methodological consistency across the field. Finally, these findings highlight the need for automated detection and removal approaches to further enhance data quality.

## Data and code availability

Data used for the statistical analyses and to generate figures are available from: https://doi.org/10.25909/32260149

Code used for analyses is available from: https://github.com/nigelrogasch/tms-eeg-step-filtering-artifacts

## Author contributions

**Marissa M. Holden:** Conceptualisation, Software, Formal analysis, Investigation, Data Curation, Writing – Original Draft, Visualisation. **Mitchell R. Goldsworthy:** Conceptualisation, Methodology, Writing - Review & Editing, Supervision. **Wei-Yeh Liao:** Investigation, Data Curation, Writing - Review & Editing. **Scott R. Clark:** Writing - Review & Editing, Supervision. **Christopher C. Cline:** Software, Writing - Review & Editing. **Corey J. Keller**: Writing - Review & Editing. **Julio C. Hernandez-Pavon:** Writing - Review & Editing. **Nigel C. Rogasch:** Conceptualisation, Methodology, Software, Validation, Formal analysis, Investigation, Data Curation, Writing – Original Draft, Visualisation, Supervision, Project Administration, Funding Acquisition.

## Supplementary materials

### Supplementary methods

#### Minimal processing pipeline 1 (for figure 1)

Data were epoched around the TMS pulse (-1000 to 1000 ms), and baseline corrected (-500 to-10 ms). The TMS pulse artifact was removed and interpolated (-2 to 10 ms; cubic interpolation), bad channels and trials were manually identified and removed, and the data were referenced to the average of all channels.

#### Minimal processing pipeline 2 (for figure 3)

First, we quantified the percentage of data points that had ‘clipped’ (i.e., moved outside the amplifier range) by counting the number of samples > |3.276| mV (i.e., the input range of the amplifier) and dividing by the total number of samples across channels. Next, we removed all the data beyond the first 30 pulses to minimise channels impacted by clipping, which increased across the duration of the recording in the DC-coupling condition (supplementary figure S1). We then removed and interpolated data around the TMS pulse in the continuous recordings (-2 to 10 ms; cubic interpolation) and resampled the data (1000 Hz). Channels showing flat lines longer than 4 s were removed, data were epoched around the TMS pulse (-1000 to 1000 ms), baseline corrected (-500 to-10 ms), and epochs with values >3 SD either across a single channel or across all channels were removed. Any missing channels were then replaced using spherical interpolation and the data were referenced to the average of all channels. Next, interpolated data and channels were removed prior to FastICA and independent components representing TMS-evoked muscle activity, blinks or eye-movement were automatically detected and removed using the default values of the *pop_tesa_compselect.m* function. Finally, removed data and channels were interpolated.

---

**Supplementary figure S1:**
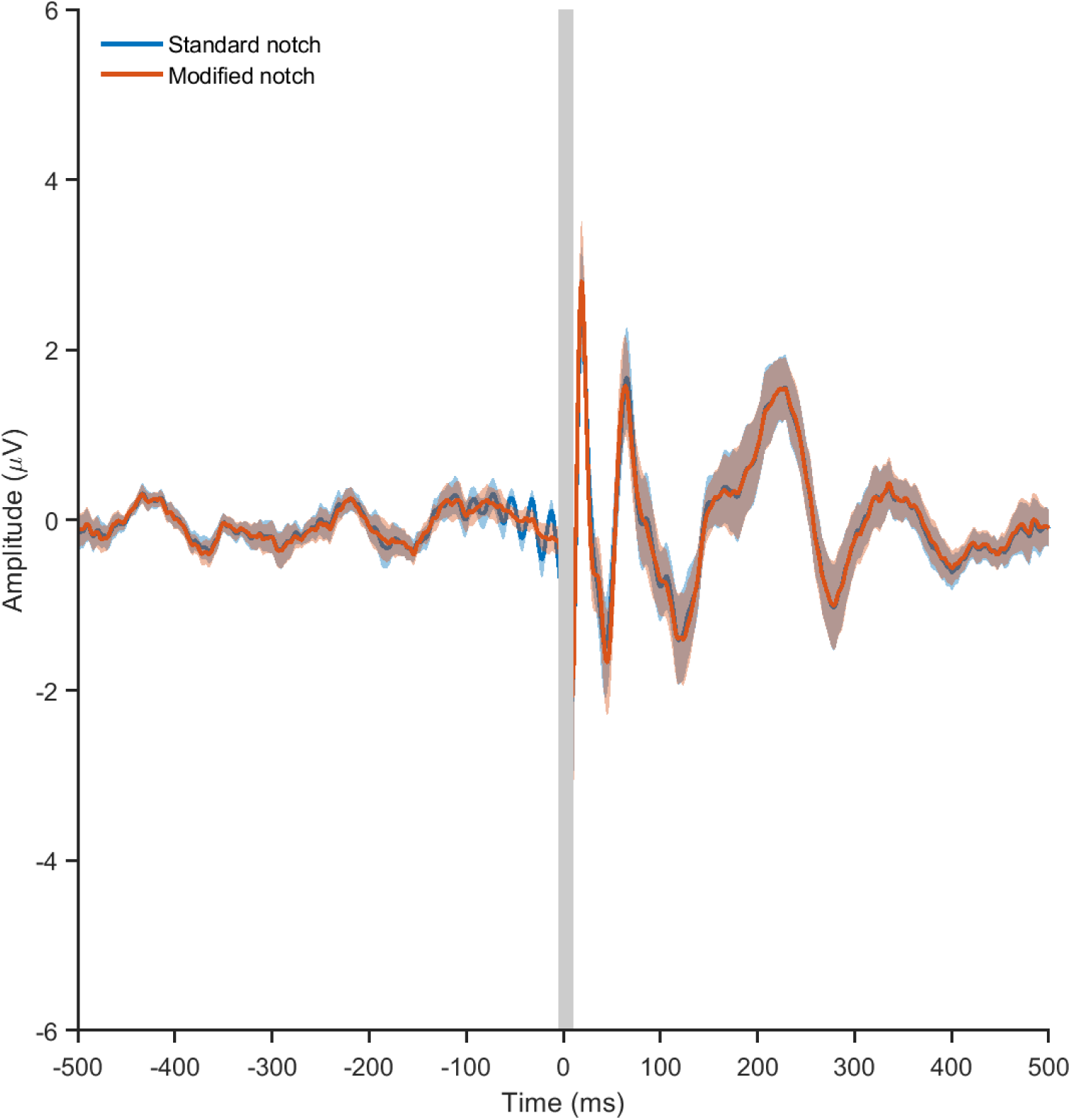
Impact of different notch filters on TEPs. Both TEPs used robust detrending and a modified high-pass Butterworth filter (1 Hz, fourth order) with autoregressive extrapolation. The blue line shows data following a standard band-stop (i.e., notch) Butterworth filter (48-52 Hz, second order). A clear ringing artifact is evident before the pulse when filtering at a frequency of ∼50 Hz. The red line show data following a modified band-stop Butterworth filter using autoregressive extrapolation (48-52 Hz, fourth order). The ringing artifact was prevented using the modified filter.

**Supplementary figure S2:**
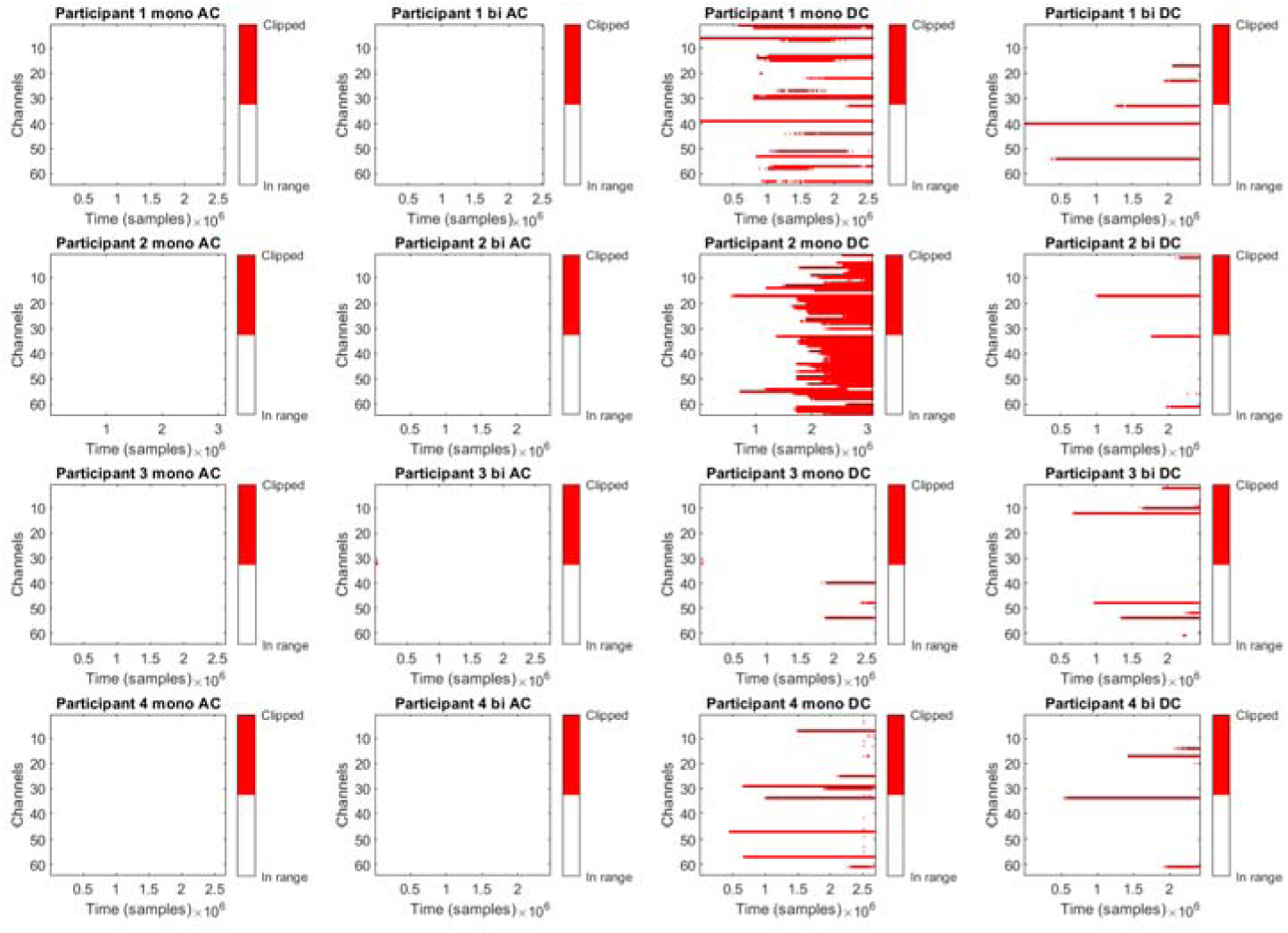
Time course of channels clipping across experimental sessions. Each panel shows a channel-by-time plot with white indicating the channel was in range, whereas red indicates the channel was clipping (e.g., >|3.676| mV). Individual participants are shown across rows whereas different conditions are shown down columns. Clipping is evident in conditions using DC-coupled amplifier settings and generally increases across the duration of the experiment. The percentage of samples lost to clipping tended to be larger following monophasic pulse shapes compared to biphasic pulses.

**Supplementary figure S3:**
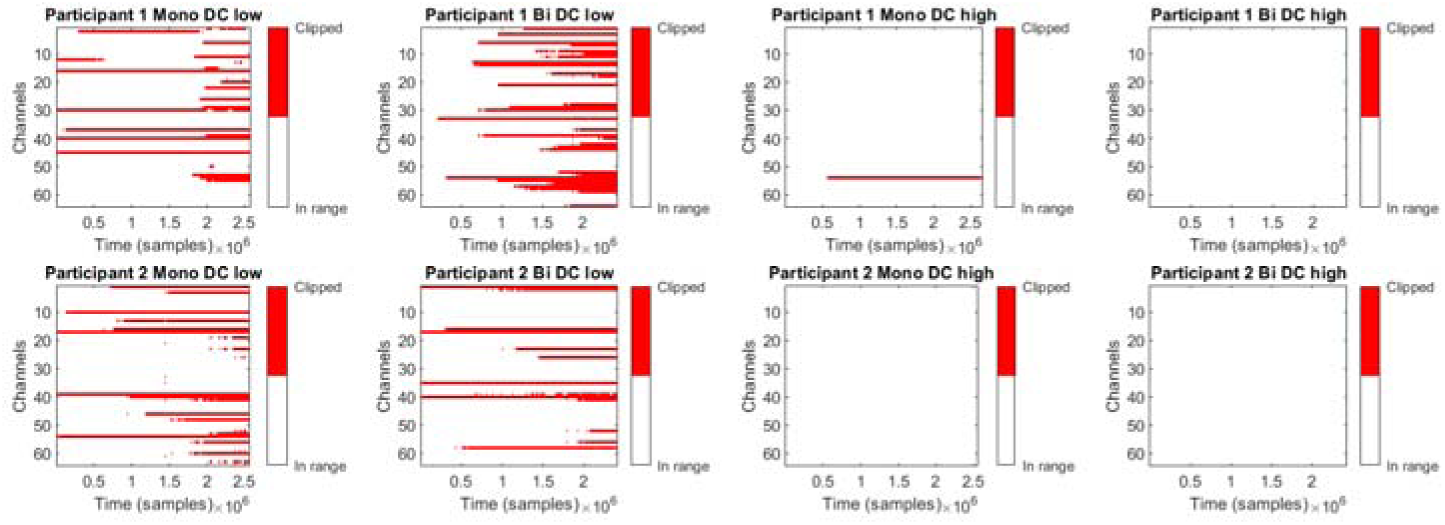
Time course of channels clipping across experimental sessions using different amplifier ranges. Each panel shows a channel-by-time plot with white indicating the channel was in range, whereas red indicates the channel was clipping (e.g., >|3.676| mV for low range; >|16.38| mV for high range). Individual participants are shown across rows whereas different conditions are shown down columns. Clipping is evident in conditions using DC-coupled amplifier settings with low amplifier range settings and generally increases across the duration of the experiment, but is decreased with high amplifier range settings.

**Supplementary figure S4:**
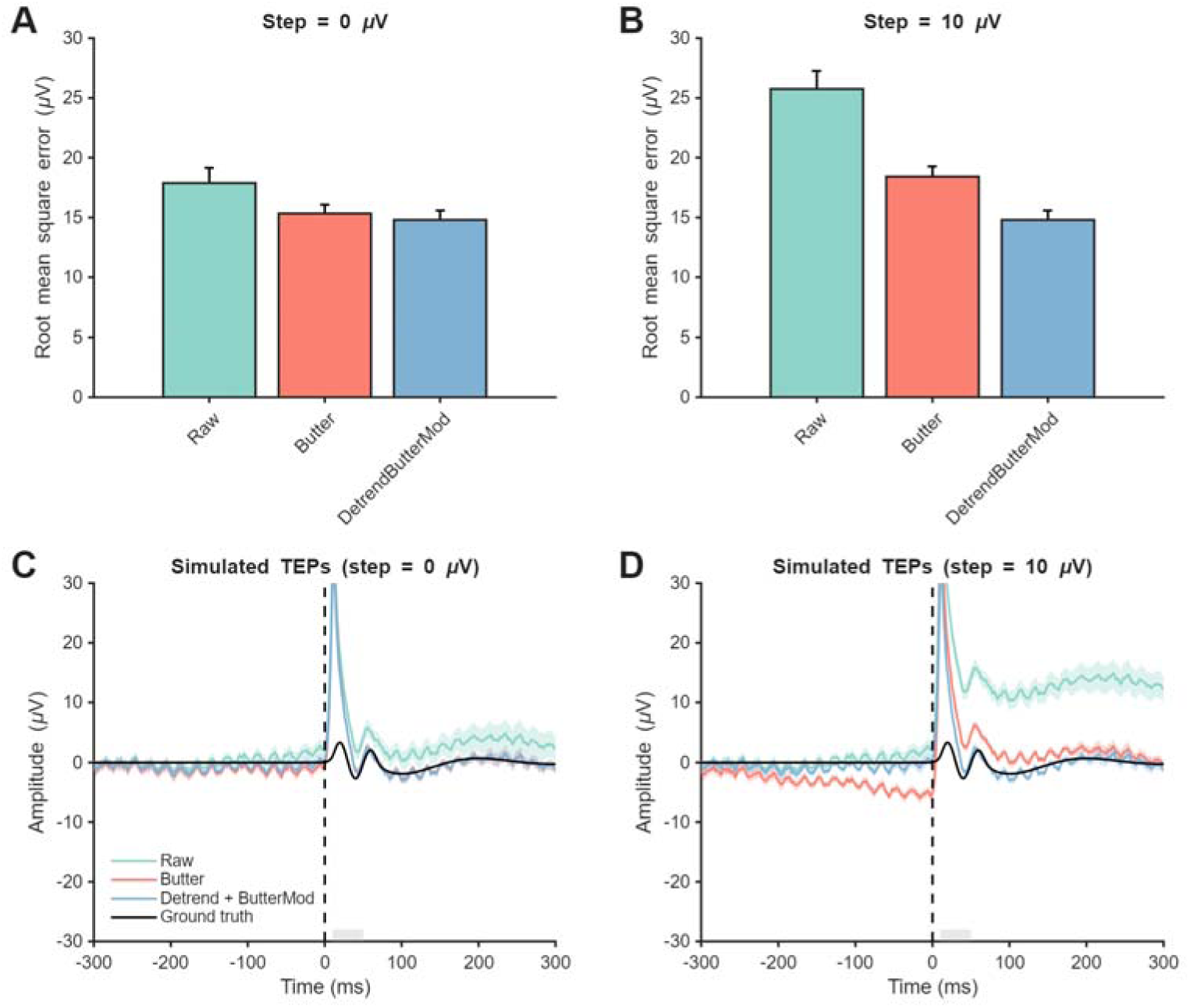
The impact of different cleaning combinations on recovering the ground truth TEP in the presence of decay/discharge artifacts. A) Root mean square error (RMSE) between the ground truth TEP and cleaned simulated TEPs embedded in real EEG data with no step artifact and a decay/discharge artifact for 2 different combinations of detrending and filtering. Bars represent the group mean and error bars the standard error of the mean (n=15). Cleaning combinations are ranked from highest to lowest RMSE. B) RMSE between the ground truth TEP and cleaned simulated TEPs embedded in real EEG data with 10 µV step artifact and decay/discharge artiact for 2 different combinations of detrending and filtering. C) Comparison of simulated TEPs embedded in real EEG data with no step artifact and decay/discharge artifact including raw data, data following Butterworth filtering, and the optimal cleaning combination (detrend + modified filter) vs. ground truth TEP. Solid lines represent group mean and shaded bars represent standard error. The grey bar indicates the time period in which RMSE was calculated. D) Comparison of simulated TEPs embedded in real EEG data with 10 µV step artifact and decay/discharge artifact including raw data, data following Butterworth filtering, and the optimal cleaning combination (detrend + modified filter) vs. ground truth TEP.

